# A biodegradable porous membrane-based lung alveoli-on-a-chip for assessing particulate-matter-induced pulmonary toxicity

**DOI:** 10.64898/2026.04.03.716404

**Authors:** Jae-Won Choi, Vaishnavi Umalkar, Xiaoliang Wang, Si-Yang Zheng

**Affiliations:** Department of Biomedical Engineering, Carnegie Mellon University, Pittsburgh, PA, USA; Division of Atmospheric Sciences, Desert Research Institute, Reno, NV, USA; Department of Electrical and Computer Engineering, Carnegie Mellon University, Pittsburgh, PA, USA

## Abstract

Understanding how airborne particulates disrupt the human alveolar barrier requires *in vitro* systems that accurately replicate its composition and function. We present a biodegradable lung alveoli-on-a-chip that reproduces the architecture and physiology of the human air-blood interface using a porous poly(lactic-co-glycolic acid) (PLGA) membrane positioned between epithelium and endothelium under air-liquid interface (ALI) culture. The membrane, fabricated by porogen-assisted nonsolvent-induced phase separation, exhibited >50 % porosity, ∼2 µm thickness, and mechanical compliance over 100-fold higher than conventional Transwell inserts, closely resembling the native interstitium. During co-culture, gradual PLGA degradation was compensated by cell-secreted extracellular-matrix (ECM) proteins such as collagen IV and laminin, forming a self-remodeling barrier that maintained integrity for at least 11 days. The platform supported stable epithelial-endothelial co-culture, high transepithelial electrical resistance, and physiologically relevant permeability. To demonstrate its utility, the chip was used to assess pulmonary toxicity of four types of waste-combustion-derived particulates, including rubber, plastic bags, plastic bottles, and textile fibers, delivered apically under ALI conditions. All combustion products reduced cell viability, increased hydrogen-peroxide release, and elevated γ-H2AX expression, indicating oxidative and genotoxic stress, while disrupting barrier permeability. Rubber combustion particles elicited the most severe toxicity, causing the greatest loss of viability, accumulation of reactive oxygen species, and formation of DNA double-strand breaks. Together, these results establish a biodegradable, ECM-remodeling lung alveoli-on-a-chip as a physiologically relevant platform for investigating source-specific particulate toxicity and alveolar-barrier pathophysiology. By bridging environmental exposure models with human-relevant lung biology, this system provides a quantitative and translatable tool for evaluating respiratory risks and therapeutic interventions.

## Introduction

The human alveolar barrier forms an expansive but microscopically thin structure that underpins efficient pulmonary gas exchange. Across an estimated surface area of roughly 100 m²,^1^ this barrier allows oxygen and carbon dioxide to diffuse rapidly while preserving essential immune protection against inhaled particulates and pathogens.^2^ Such efficiency originates from its refined microanatomy: a single epithelial layer lies adjacent to a capillary endothelium, the two compartments divided only by a nanoscale interstitial region containing basement membranes, extracellular matrix components, and supporting stromal cells. Disturbance of this fragile arrangement is a hallmark of numerous respiratory disorders—including viral and bacterial pneumonia, chronic obstructive pulmonary disease (COPD), acute respiratory distress syndrome (ARDS), pulmonary fibrosis, and asthma—as well as exposure-related injuries from airborne particulates or cigarette smoke.^3–5^ Collectively, respiratory diseases impose a profound global health and socioeconomic burden, with COPD, lower respiratory infections, and lung cancer persistently listed among the world’s ten most common causes of death.^6^ Faithfully replicating this alveolar interface *in vitro* is therefore essential for advancing the understanding of pulmonary pathophysiology and developing effective therapeutics.

While animal models provide valuable whole-organism context for studying pulmonary function and therapy, their interspecies variability and limited control over alveolar exposure make quantitative interpretation challenging.^7^ Although two-dimensional culture formats are easy to handle and widely scalable, they cannot replicate the spatial organization or mechanical cues characteristic of native lung tissue.^8^ While Transwell® systems enable air–liquid interface culture, their dense, non-interconnected membranes substantially hinder molecular diffusion and intercellular communication across the barrier.^9^ To overcome these limitations, organ-on-a-chip technologies have emerged as powerful tools to reproduce organ-level physiology and disease mechanisms *in vitro*.^10,11^

Despite these advances, a key physiological constraint of most lung-on-a-chip devices lies in the barrier membrane. In the native alveolus, the septal barrier possesses a remarkably high porosity (close to 80%) and enables gas diffusion through submicron-scale pathways, whereas most lung-on-a-chip membranes remain relatively dense, with less than 10% open area, poorly connected pores, and thickness several times greater than that of the natural interstitium.^12–18^ Moreover, most synthetic membranes are mechanically stiff and non-degradable, making it difficult to study dynamic remodeling or fibrosis. These structural limitations hinder oxygen transport and disrupt both intercellular and soluble-factor signaling across the interface, reducing the fidelity of epithelial–endothelial crosstalk.

Here, we address these limitations by engineering a lung alveoli-on-a-chip featuring a biodegradable poly(lactic-co-glycolic acid) (PLGA) membrane that serves as an interstitium-like scaffold positioned between the apical epithelium and basolateral endothelium. The PLGA membrane exhibited a porosity exceeding 50% and a thickness of approximately 2 µm, making it 4.6-fold thinner, 5.9-fold more porous, and 114.9-fold more compliant than the Transwell® insert, thus more closely approximating the physical and mechanical characteristics of the native alveolar barrier.

Open burning of solid waste is a widespread practice in many regions, particularly in developing countries lacking adequate waste management infrastructure and in military operations utilizing open-air burn pits for waste disposal.^19–22^ Such uncontrolled combustion of waste releases a complex mixture of gases and fine particulate matter (PM), contributing substantially to both local and global air pollution. Waste combustion products, which contain environmentally persistent free radicals (EPFRs), transition metals, polycyclic aromatic hydrocarbons (PAHs), and chlorinated organics, are known to generate reactive oxygen and chlorine species in the atmosphere.^23,24^ These reactions not only degrade air quality but also promote secondary aerosol formation and oxidative atmospheric chemistry, ultimately exacerbating climate forcing and human exposure risks.^25,26^

Growing evidence indicates that exposure to waste-combustion-derived particulates poses significant health hazards. Epidemiological and toxicological studies have linked these emissions to oxidative stress, genotoxicity, and systemic inflammation, which mainly contribute to cardiovascular and respiratory diseases.^27,28^ Populations residing near open-burning sites or engaged in informal waste collection are at particularly high risk due to prolonged exposure to toxic combustion emissions.^29^ Despite these concerns, most existing assessments rely on environmental monitoring or *in vitro* cytotoxicity assays using single cell types,^30–32^ which fail to reproduce the complex structure and microenvironment of the human lung. Consequently, the mechanisms by which inhaled waste- combustion particulates interact with the alveolar barrier, affecting epithelial–endothelial integrity, oxidative balance, and DNA stability, remain largely unexplored.

As a proof-of-concept application, we employed the developed lung alveoli-on-a-chip to directly expose epithelial–endothelial co-cultures to four distinct types of waste-combustion products. Subsequent analyses quantified cell death, barrier disruption, oxidative stress, and DNA damage. Among the tested materials, rubber-combustion particulates induced the most severe pulmonary toxicity. These findings demonstrate that this biodegradable lung-on-a-chip serves as a physiologically relevant platform for investigating PM-induced lung injury, with direct implications for occupational exposure assessment and public health policy.

### Design and fabrication of the lung alveoli-on-a-chip with porous PLGA membranes

We developed a lung alveoli-on-a-chip that recapitulates the microenvironment of the human alveolus. *In vivo*, the air–blood barrier consists of an alveolar epithelial layer and a capillary endothelial layer separated by an ultrathin interstitial space containing a continuous basement membrane typically 1∼2 µm thick (Fig. 1a).^33^ To emulate this physiological architecture, we fabricated biodegradable porous PLGA membranes that serve as the core structural scaffold of the lung alveoli-on-a-chip.

**Figure 1.**
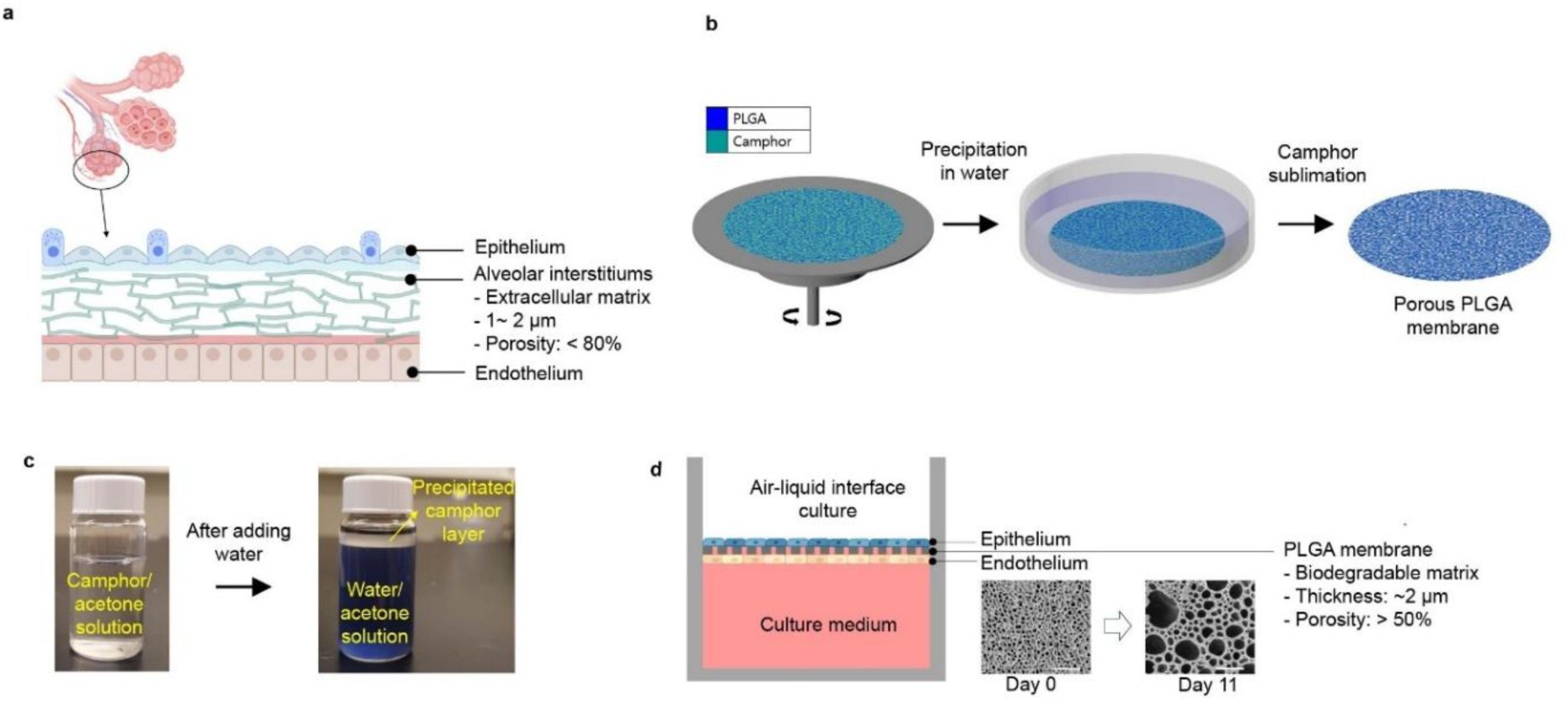
Design of the lung alveoli-on-a-chip with biodegradable porous PLGA membranes. **a,** Schematic illustration of the lung alveolar barrier structure. The alveolar wall consists of an epithelial layer, an endothelial layer, and an intervening alveolar interstitium. **b,** Schematic illustration of the fabrication process of porous PLGA membranes. **c,** Optical images of the camphor-acetone solution (left) and the precipitated camphor layer after mixing of the camphor-acetone solution in colored water (right). **d,** Schematic illustration of the lung alveoli-on-a-chip system under air–liquid interface (ALI) culture, showing epithelial and endothelial cell layers separated by a biodegradable PLGA membrane. Representative SEM images of the PLGA membrane at Day 0 and Day 11 demonstrate increased porosity during degradation. Scale bars: 10 µm.

We selected PLGA as the membrane material because it is biodegradable and biocompatible,^34^ making it well suited for constructing physiologically relevant tissue models. An interconnected porous architecture was engineered by integrating nonsolvent-induced phase separation (NIPS) with camphor-templated dendritic crystallization, in which camphor transiently served as a pore-forming agent during polymer solidification.^35^ In brief, a PLGA–camphor solution prepared in acetone was deposited onto the substrate by spin coating (Fig. 1b). Immersing the film in water, which is a nonsolvent for both PLGA and camphor, triggers a rapid solvent exchange as acetone diffuses outward while water penetrates the polymer matrix. This process induced PLGA phase separation and solidification, while camphor simultaneously crystallized into a dendritic framework within the polymer matrix (Fig. 1c).^36^ Following freeze-drying, camphor was removed via sublimation, leaving behind interconnected voids that replicated its crystalline morphology.

PLGA membranes were engineered to faithfully recapitulate the structural and functional characteristics of the native interstitium and were positioned between the apical and basolateral chambers molded in polydimethylsiloxane (PDMS) (Fig. 1d). This setup allowed the establishment of an air–liquid interface (ALI) culture, in which epithelial cells were cultivated on the apical side of the membrane, while endothelial cells were grown on the basolateral side. During co-culture within the chip, the PLGA membrane gradually degraded, exhibiting a porosity exceeding 50% and a reduced thickness of approximately 2 µm after 11 days.

### Degradation behavior and permeability of porous PLGA membranes compared with Transwell® inserts

To fully characterize the hydrolytic stability of the PLGA membrane under biocompatible aqueous conditions, samples were incubated in phosphate-buffered saline (PBS) at 37 °C for 11 days to monitor morphological and functional changes over time. Spin-coated porous PLGA membranes exhibited a dual-sided asymmetric structure, with smaller pores on the apical surface and larger pores on the basolateral side. Initially, the apical surface showed 20.5% porosity with a mean pore size of 0.94 ± 0.26 µm, while the basolateral surface exhibited 25.5% porosity with a mean pore size of 1.11 ± 0.48 µm (Fig. 2a–c). As the membrane underwent hydrolytic degradation in PBS at 37 °C, its porosity gradually increased. By day 11, the apical surface porosity increased to 53.5% with a mean pore size of 3.79 ± 2.11 µm, and basolateral surface porosity reached 55.8% with a mean pore size of 3.84 ± 2.10 µm. After 11 days of degradation, the apical and basolateral porosities of PLGA membranes were approximately 5.6-fold and 5.9-fold higher than those of 0.4 µm pore-sized Transwell® membrane commonly used in lung-on-a-chips, respectively (Fig. S1). Before degradation, the PLGA membrane had a thickness of 5.85 µm, which decreased to 2.33 µm after 11 days. The final thickness was approximately 4.6-fold thinner than the Transwell® insert and closely resembled the ultrathin septal layers of the native alveolar barrier *in vivo* (Fig. 2d). The membrane readily allowed aqueous solutions to pass through its thickness, as evidenced by the rapid penetration of a water-soluble dye, indicating a highly permeable and hydrophilic pore network well suited for ALI applications (Fig. 2e). Mass-loss analysis further corroborated the progressive thinning and increased surface porosity of PLGA membranes during degradation. After 15 days in PBS, 52.8% of the initial mass was lost, corresponding to an average degradation rate of approximately 3.5% per day, consistent with hydrolytic degradation (Fig. 2f). Membrane permeability, a critical indicator of intercellular crosstalk and nutrients, growth factors, and signaling molecules, was substantially enhanced compared to the Transwell® membrane. Before degradation, the apparent permeability of PLGA membranes was 3.2-fold higher, and after 11 days of degradation, it increased to 9.1-fold higher than that of Transwell® membrane (Fig. 2g).

**Figure 2.**
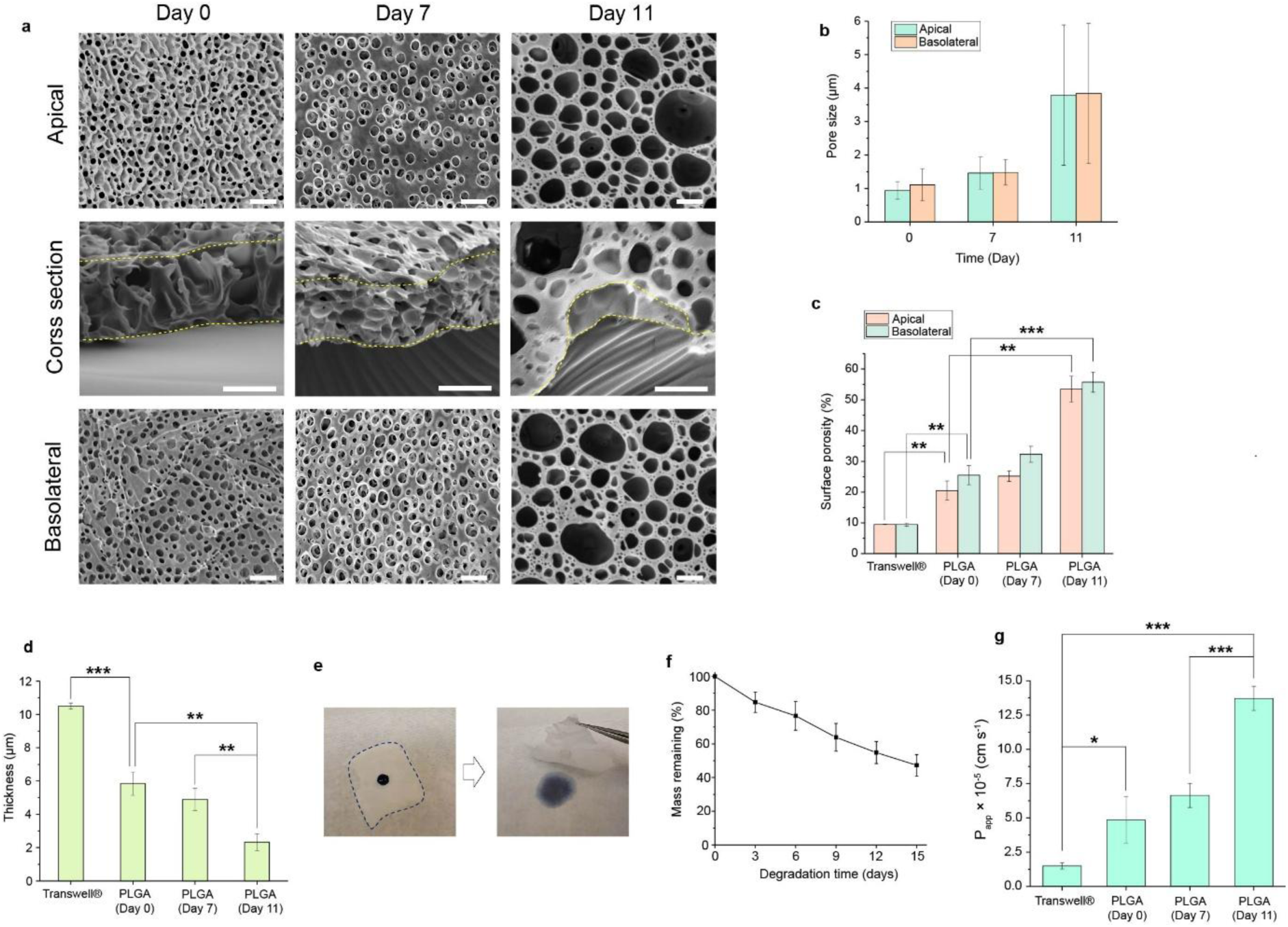
Degradation characteristics of PLGA membranes. **a,** Representative SEM images of PLGA membranes after degradation in PBS at 37 °C for 0, 7, and 11 days, illustrating the apical, cross-sectional, and basolateral morphologies. Scale bar: 5 µm. **b,** Changes in pore size on the apical and basolateral surfaces of PLGA membranes during degradation in PBS at 37 °C for 0, 7, and 11 days. **c,** Surface porosity of the apical and basolateral sides of Transwell® membranes and PLGA membranes after degradation in PBS at 37 °C for 0, 7, and 11 days. **d,** Thickness of Transwell® membranes and PLGA membranes after degradation in PBS at 37 °C for 0, 7, and 11 days (n = 4). **e,** Optical images of a drop of water-soluble ink on the PLGA membrane (dashed lines indicate the contour of the PLGA membrane) before and after the ink has passed through the membrane. **f,** Mass loss of porous PLGA membranes during degradation in PBS at 37 °C over time (n = 4). **g,** Apparent permeability (P_app_) of 4 kDa dextran across Transwell® membranes and PLGA membranes after degradation in PBS at 37 °C for 0, 7, and 11 days (n = 4).

### Physicochemical characterization of porous PLGA membranes

To assess the mechanical integrity of the PLGA membrane, the apparent tensile modulus of each membrane was determined from its stress–strain curves and compared with that of Transwell® insert (Fig. 3a-b, Fig. S2-S3). Before degradation, the PLGA membrane exhibited an apparent Young’s modulus of 11.59 MPa, which was approximately 34.7-fold lower than that of the Transwell®,^37^ indicating its higher flexibility and compliance. As the PLGA membrane underwent hydrolytic degradation over 11 days, the modulus further decreased to 3.50 MPa. The gradual reduction in tensile modulus was accompanied by a decrease in strain at break and ultimate tensile strength (UTS), which dropped by 9.0-fold and 4.0-fold, respectively, consistent with the progressive increase in porosity observed during degradation (Fig. 3c–d). Fourier transform infrared (FT-IR) spectra of the porous PLGA membrane revealed a distinct carbonyl stretching vibration at 1750 cm⁻¹, while the camphor-associated ketone band at 1743 cm⁻¹ was absent, confirming that no residual camphor remained after fabrication (Fig. 3e). The pristine PLGA surface showed a water contact angle of 90.8°, indicating moderate hydrophobicity. After gelatin coating applied prior to cell seeding, the contact angle was reduced to 64.3°, demonstrating enhanced surface wettability favorable for subsequent cell attachment (Fig. 3f).

**Figure 3.**
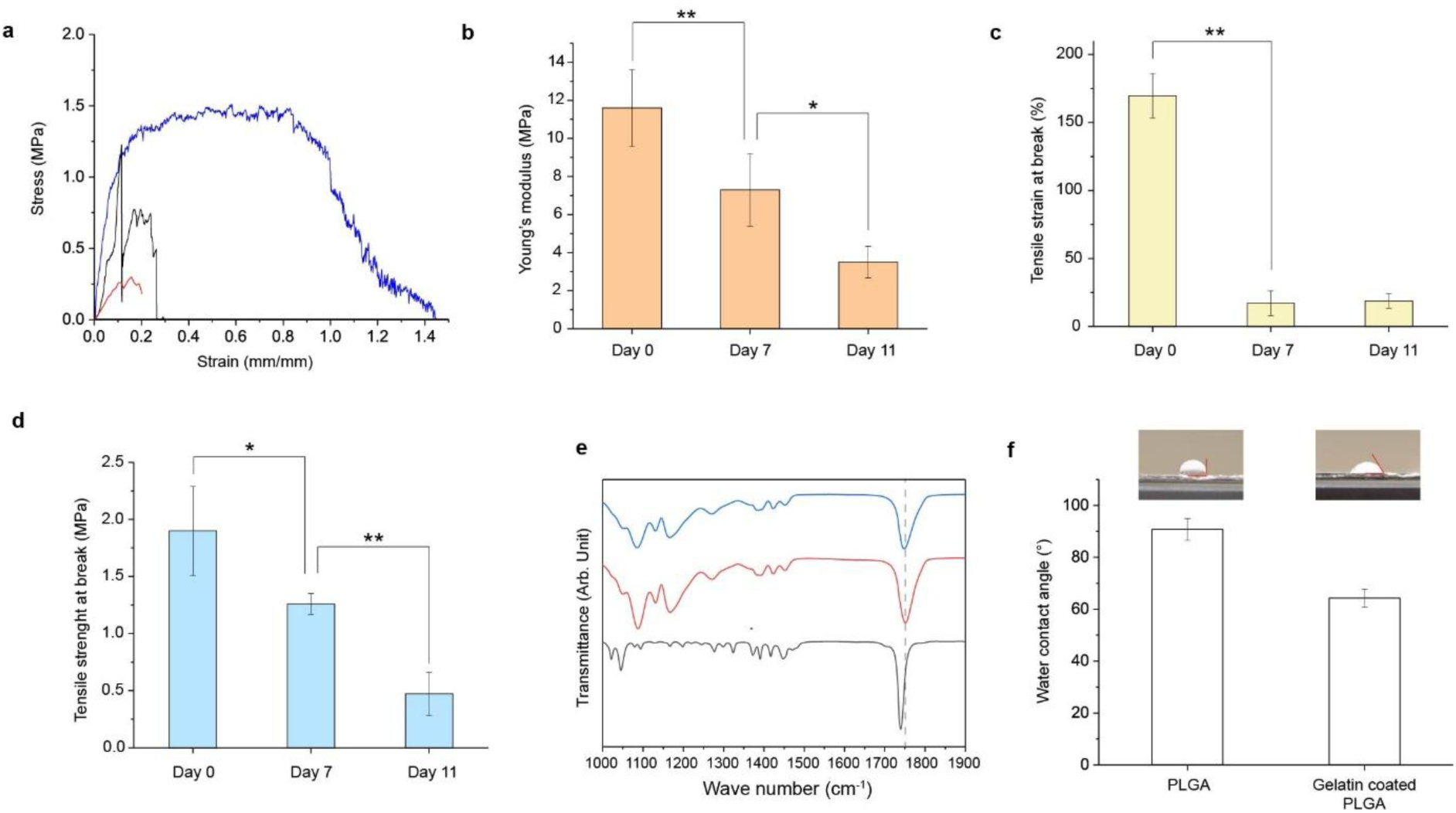
Physicochemical characteristics of PLGA membranes. **a,** Representative stress–strain curves of PLGA membranes after degradation in PBS at 37 °C for 0 (blue), 7 (black), and 11 days (red), obtained from tensile strength tests. **b,** Young’s modulus of PLGA membranes after degradation in PBS at 37 °C for 0, 7, and 11 days (n = 4). **c,** Tensile strain at break of PLGA membranes after degradation in PBS at 37 °C for 0, 7, and 11 days (n = 4). **d,** Ultimate tensile strength of PLGA membranes after degradation in PBS at 37 °C for 0, 7, and 11 days (n = 4). **e,** FT-IR spectra of camphor (black), PLGA membrane (red), and pure PLGA (blue). **f,** Water contact angle measurements of bare PLGA membranes and gelatin-coated PLGA membranes (n = 4).

### Establishment of epithelial and endothelial ALI co-culture on lung alveoli-on-a-chip

To fabricate the lung alveoli-on-a-chip device, the PLGA membrane was placed between the PDMS apical and basolateral chambers and bonded using uncured PDMS as an adhesive layer. Human primary alveolar epithelial cells (HPAECs) and human lung microvascular endothelial cells (HMVECs) were established on opposing sides of the PLGA membrane under ALI culture, forming a barrier architecture analogous to the native alveolus. To establish the epithelial–endothelial co-culture, endothelial cells were first introduced onto the basolateral surface of the inverted device. After an incubation of 3 hours to facilitate firm attachment, the chip was repositioned upright, and epithelial cells were then deposited onto the apical surface. Liquid–liquid culture was maintained for seven days prior to transition to ALI culture. After 11 days of on-chip culture, the viability of both HPAECs and HMVECs exceeded 95% (Fig. 4a–b). Under ALI conditions, cells formed confluent monolayers on the PLGA membrane. Tight junctions in HPAECs were visualized by ZO-1 immunostaining, whereas adherens junctions in HMVECs were identified by VE-cadherin staining (Fig. 4c–d). Cross-sectional SEM imaging demonstrated well-defined epithelial and endothelial layers cultured on opposite sides of the PLGA membrane (Fig. 4e). Remarkably, the interlayer distance between the two cell types was comparable to that found in native alveoli. Barrier integrity was evaluated by measuring transepithelial/transendothelial electrical resistance (TEER). Under ALI conditions, HPAECs exhibited a TEER value of 336.1 Ω·cm², which increased to 372.3 Ω·cm² upon co-culture with HMVECs (Fig. 4f). Under ALI conditions, the permeability of the HPAEC–HMVEC co-culture decreased to 1.99 ± 0.38 × 10⁻⁵ cm·s⁻¹, representing a 6.8-fold reduction relative to cell-free membranes (Fig. 4g). This indicates that the system successfully recapitulates the physiological roles of the alveolar interface, where epithelial layers provide a protective barrier and the capillary endothelium regulates fluid exchange within the lung microenvironment.^39,40^ The highly porous and interconnected architecture of the PLGA membrane promoted efficient gas and nutrient exchange and enabled bidirectional paracrine signaling.^41–44^ The chip platform retained both structural stability and functional integrity even when human umbilical vein endothelial cells (HUVECs) were used instead of HMVECs (Fig. S4–S8).

**Figure 4.**
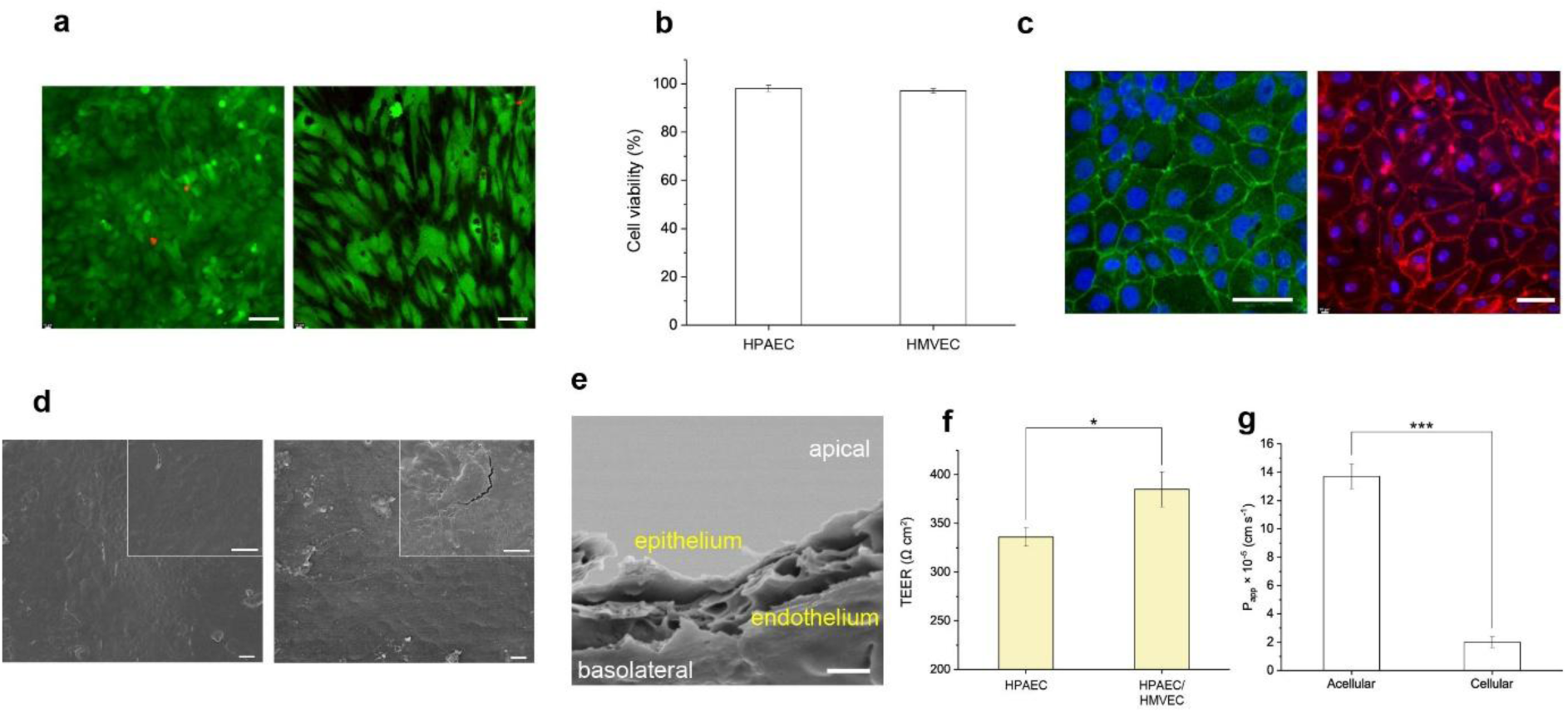
Establishment of epithelial and endothelial ALI co-culture on lung alveoli-on-a-chip. **a,** Live/dead staining images of co-cultured cells in the chip after 11 days of culture: HPAECs (left) and HMVECs (right). Live cells are shown in green and dead cells in red. Scale bars: 50 µm. **b,** Cell viability in the chip after 11 days of HPAEC–HMVEC co-culture. **c,** Representative immunofluorescence images of cells co-cultured on PLGA membranes: HPAECs stained for ZO-1 (left) and HMVECs stained for VE-cadherin (right). Nuclei are counterstained with DAPI (blue). Scale bars: 50 µm. **d,** Representative top-view SEM images of cells cultured on PLGA membranes after 11 days of co-culture: HPAECs (left) and HMVECs (right). Insets show higher-magnification images. Scale bars: 10 µm. **e,** Representative cross-sectional SEM image of cells cultured on porous PLGA membranes under HPAEC–HMVEC co-culture conditions. The medium in the apical chamber was removed to expose the epithelial cells to air, establishing an ALI culture. Scale bar: 2 µm. **f,** Transepithelial electrical resistance (TEER) in the lung alveoli-on-a-chip for HPAEC monoculture and HPAEC–HMVEC co-culture (n = 4). **g**, Apparent permeability (P_app_) of 4 kDa dextran across PLGA membranes under two conditions: acellular (no cells) and cellular (co-culture of HPAECs and HMVECs maintained for 11 days on PLGA membranes) (n = 4).

### ECM formation compensating for membrane degradation in the lung alveoli-on-a-chip

To investigate whether extracellular matrix (ECM) formation compensates for the progressive degradation of PLGA membranes during co-culture, we monitored ECM deposition and membrane integrity over time using FITC-labeled PLGA membranes (Fig. 5a, Fig. S9). Immunofluorescence imaging revealed that both collagen IV and laminin, the major structural components of the alveolar basement membrane,^45^ were increasingly deposited by co-cultured HPAECs and HMVECs from day 1 to day 11 (Fig. 5b-c). Concurrently, the fluorescence intensity from the FITC-labeled PLGA membrane decreased, indicating substantial degradation, whereas the intensities of collagen IV and laminin signals significantly increased (Fig. 5d). These results suggest that ECM proteins secreted by epithelial and endothelial cells progressively replaced the degrading PLGA scaffold, thereby recapitulating the microenvironment of native lung tissue that is essential for maintaining barrier function and mediating cell signaling.

**Figure 5.**
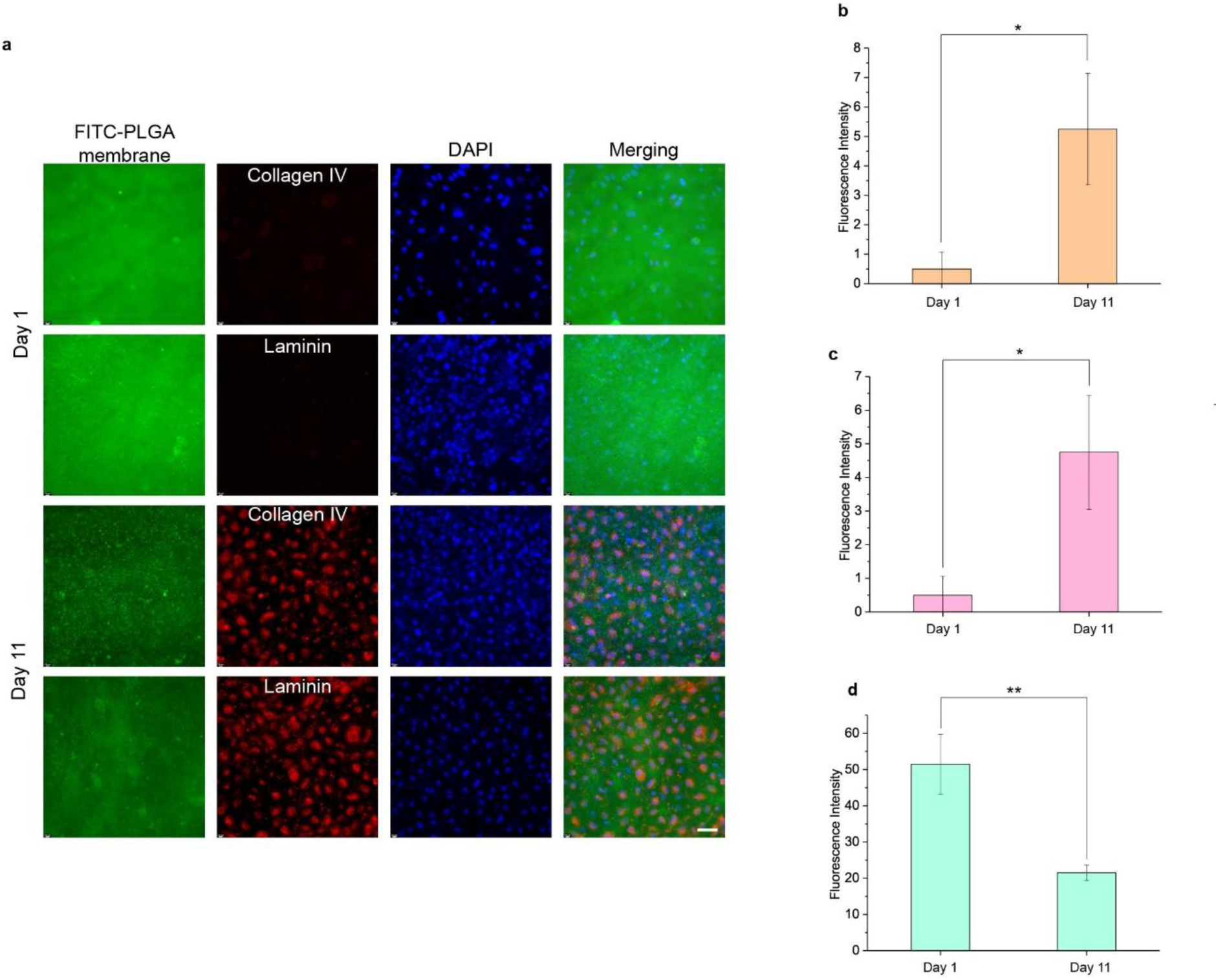
ECM formation compensating for membrane degradation in the lung alveoli-on-a-chip. **a** Representative immunofluorescence images of co-cultured HPAECs and HMVECs on FITC-labeled PLGA membranes (green) at day 1 and day 11, stained for collagen IV and laminin (red). Scale bar: 50 µm. **b,** Fluorescence intensity of laminin after 1 and 11 days of cell culture in the chip system (n = 4). **c,** Fluorescence intensity of collagen IV after 1 and 11 days of cell culture in the chip system (n = 4). **d,** Fluorescence intensity of FITC-labeled PLGA membranes after 1 and 11 days of cell culture (n = 4).

### Toxicological effects of waste combustion products on lung cells cultured in the lung alveoli-on-a-chip

Airborne PM is recognized as the fifth leading cause of mortality worldwide and a major driver of cardiopulmonary morbidity.^46,47^ However, the toxicological impact of PM varies considerably depending on its chemical composition and emission source.^48^ Understanding these source-specific toxicities is critical for developing effective, health-oriented air-quality policies and mitigation strategies. Among the diverse PM sources, waste combustion products represent a complex and underexplored class of pollutants. Uncontrolled open burning of household and municipal waste releases fine particulates enriched with transition metals, polycyclic aromatic hydrocarbons (PAHs), and environmentally persistent free radicals (EPFRs), all of which can trigger oxidative stress and genotoxic damage upon inhalation.^23,24^ Despite their ubiquity in ambient air, the direct pulmonary consequences of exposure to waste-derived particulates remain poorly defined, largely due to the lack of physiologically relevant *in vitro* models capable of recapitulating the alveolar interface.

To evaluate the pulmonary toxicity of combustion-derived particulates, PM_2.5_ (particles with aerodynamic diameter ≤ 2.5 µm) emitted from burning of four representative waste materials (rubber, plastic bags, plastic bottles, and textile) were directly delivered to the apical surface of the lung alveoli- on-a-chip under ALI conditions at the same dosage of 50 µg/cm² (Fig. 6a). Although the actual airborne concentrations of combustion-derived particulates are generally limited to the range of 0.1–9 × 10⁻⁴ µg/cm³, a higher dosage was applied in this study to facilitate observable toxicological effects within a limited experimental timeframe.^49–51^ Exposure to these particulates markedly reduced cell viability across epithelial–endothelial co-cultures after 24 hours (Fig. 6b–c). Among the tested materials, rubber combustion particulates caused the most severe cytotoxicity, decreasing HPAEC and HMVEC viability to 52.5% and 57.9%, respectively. The comparable reduction in epithelial and endothelial viability suggests that these particulates can readily traverse the epithelial layer and porous membrane to affect the capillary compartment.

**Figure 6.**
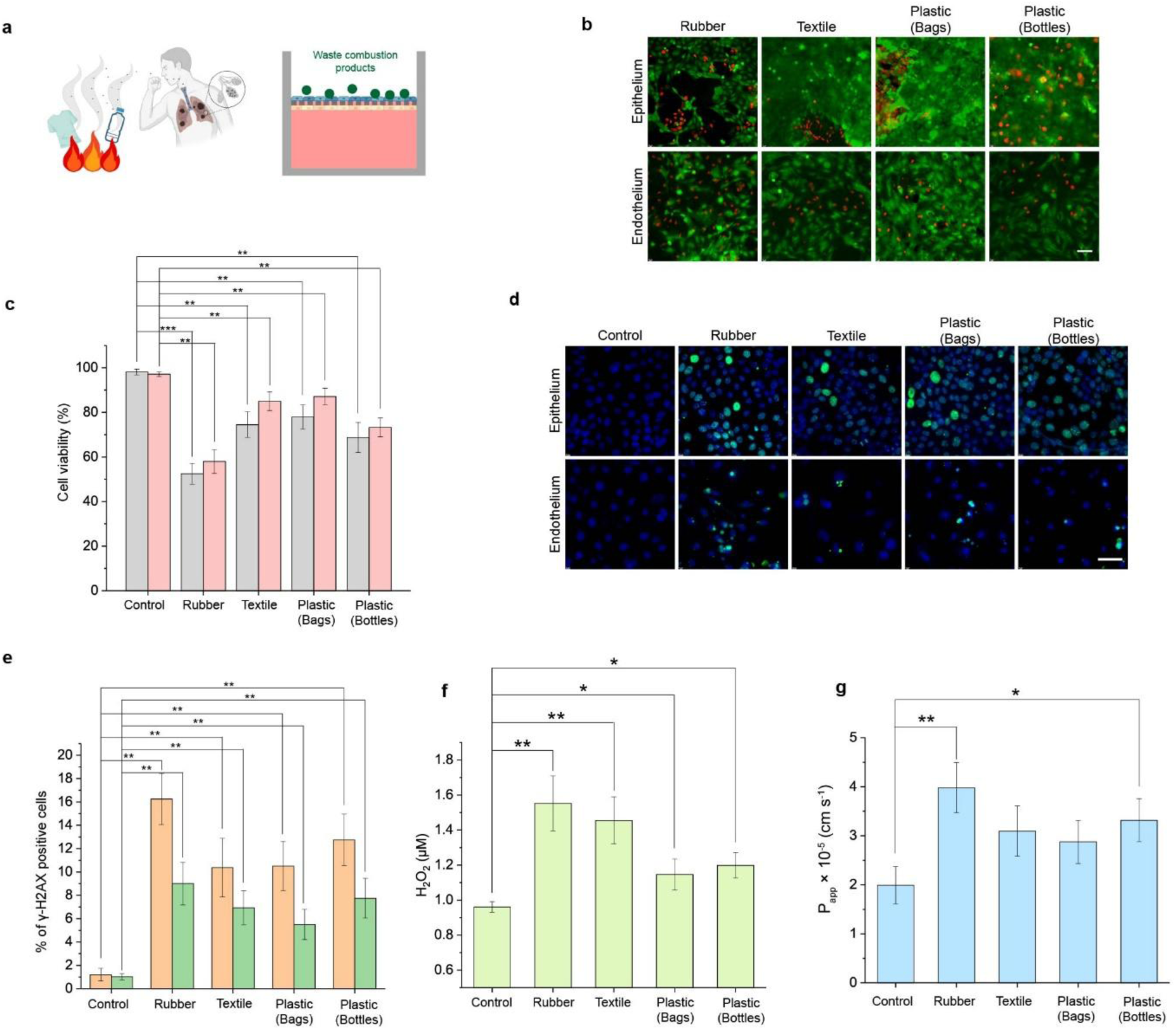
Toxicological effects of waste combustion products on the lung alveoli-on-a-chip system. **a** Schematic diagram illustrating the toxicity assessment of waste combustion products using the lung alveoli-on-a-chip system. Under ALI condition, waste combustion products (50 µg/cm²) were introduced into the apical chamber, and the ALI culture was maintained for 24 hours. **b,** Live/dead staining images of epithelial cells (top row) and endothelial cells (bottom row) in the lung alveoli-on-a-chip after 24-hour exposure to four different waste combustion products. Scale bars: 50 µm. **c,** Quantification of cell viability of epithelial (gray) and endothelial (pink) cells in the lung alveoli-on-a-chip after 24-hour exposure to four different waste combustion products (n = 4). **d,** Immunostaining of γ-H2AX showing DNA double-strand breaks in epithelial and endothelial cells cultured in the lung alveoli-on-chip system after 24 hours of exposure to four different waste combustion products. Scale bar: 50 µm. **e,** Percentage of γ-H2AX–positive cells in the lung alveoli-on-chip system after 24 hours of exposure to four different waste combustion products (orange: epithelial cells, green: endothelial cells; n = 4). **f,** Reactive oxygen species (ROS) levels after treatment with four different waste combustion products. Hydrogen peroxide (H₂O₂) concentration was quantified 24 hours after exposure (n = 4). **g,** Apparent permeability (P_app_) of the lung alveoli-on-chip system to FITC–dextran (4 kDa) after 24 hours of exposure to four different waste combustion products under ALI condition (n = 4).

DNA double-strand breaks, assessed by the phosphorylated form of histone H2AX (γ-H2AX), serve as a hallmark of genotoxic damage. In the absence of exposure, the γ-H2AX–positive fraction was minimal (HPAEC: 1.2%, HMVEC: 1.0%), representing the baseline level of DNA damage. After exposure to all four types of waste combustion products, the proportion of γ-H2AX–positive cells increased markedly to 5.5–16.3% in both epithelial and endothelial populations, depending on the origin of the particulates (Fig. 6d-e). Among them, rubber combustion particulates induced the most pronounced genotoxic damage. The observed increase in γ-H2AX expression demonstrates that waste combustion products induce DNA double-strand breaks across epithelial–endothelial co-cultures.

We next examined whether exposure to waste combustion products induces reactive oxygen species (ROS) generation. All four combustion products increased hydrogen peroxide release, particularly in chips exposed to rubber particulates, indicating a markedly enhanced oxidative burden (Fig. 6f).

All four types of waste combustion products disrupted the epithelial–endothelial layers, leading to increased permeability, with rubber combustion particles causing the most pronounced effect (Fig. 6g). This trend is consistent with the observed decreases in cell viability and the increases in DNA damage and ROS generation.

Collectively, our results indicate that combustion products differ markedly in their pulmonary toxicity profiles, with rubber-derived particulates eliciting the greatest cellular damage, oxidative stress, and barrier dysfunction.

## Conclusion and Discussion

We developed a biodegradable lung alveoli-on-a-chip that reproduces the essential architecture and function of the human alveolar interface through a porous PLGA membrane acting as an interstitium-like scaffold. The membrane exhibited high open porosity (>50%) and ultrathin thickness (∼2 µm), enabling efficient gas and nutrient transport while supporting co-culture of epithelial and endothelial cells under ALI conditions. During degradation, progressive loss of PLGA mass was compensated by cell-secreted ECM proteins, including collagen IV and laminin, which gradually replaced the scaffold and preserved barrier integrity—thereby recapitulating native alveolar remodeling.

Compared with conventional Transwell® systems, this lung alveoli-on-a-chip demonstrated markedly improved permeability and physiological fidelity, effectively recreating a microenvironment that closely mirrors the *in vivo* alveolar barrier. The biodegradable PLGA-based platform supports co-culture and promotes matrix remodeling, thereby bridging the gap between traditional *in vitro* assays and physiologically relevant lung-on-a-chip models.

To demonstrate its utility, the chip was applied to investigate the pulmonary toxicity of waste combustion products, a major but underexplored source of airborne particulate pollution. Direct apical exposure to PM_2.5_ emitted from combustion of four representative type of waste materials (rubber, plastic bags, plastic bottles, and textile) revealed source-specific toxicological profiles. Previously, we characterized the chemical compositions of these combustion products in detail, showing that the particulates are acidic.^21^ The key toxic compounds relevant to biological responses are summarized in Table S1. Hydrogen chloride (HCl), a hazardous chemical known to induce acute pulmonary injury upon inhalation,^52,53^ was emitted at the highest level from rubber combustion, being 6.1 times higher than that from textile, the second-highest source. Even though HCl is in gas phase, the high elemental level of particulate sulfur (S) and chlorine (Cl), in addition to their water-soluble ions, indicate the presence of potentially toxic S- and Cl-compounds. Heavy metals (e.g., Cu, Zn, Cd, Sb, Ce, and Pb), which have been reported to cause cellular damage, apoptosis, oxidative stress, and DNA damage,^54,55^ were also most abundantly released from rubber combustion. Phthalates, PAHs, and nitro-PAHs—known for their carcinogenicity and oxidative stress-inducing potential—were either most or the second most abundant in rubber emissions.^56–62^ When considering the overall abundances of the four toxic chemical classes (Cl- and S-compounds, heavy metals, PAHs, and phthalates) generated from fresh smoke, rubber combustion produced the highest levels of hazardous compounds. This high emission burden is likely responsible for the pronounced cytotoxic effects observed with rubber particulates, which exhibited the strongest cellular damage characterized as decreased viability, elevated ROS generation, increased γ-H2AX expression indicating DNA double-strand breaks, and disruption of the epithelial–endothelial barrier function. These results highlight the ability of the lung alveoli-on-a-chip platform to quantitatively assess oxidative, genotoxic, and barrier responses within a physiologically relevant human model.

This study establishes a versatile and translatable tool for investigating the interactions between environmental particulates and the alveolar barrier. The combination of biodegradable membrane architecture, ECM remodeling, and human-relevant co-culture allows systematic exploration of pulmonary responses to PM exposures. Future development integrating immune or stromal components and embedded sensors for real-time readouts will further enhance its utility for studying inflammation, fibrosis, and regeneration. Overall, this biodegradable lung alveoli-on-a-chip represents a step forward in organ-on-chip engineering, advancing from static membrane assays toward a self-remodeling, physiologically informed system that bridges environmental toxicology, pulmonary pathophysiology, and therapeutic discovery.

## Material and Methods

### PLGA Membrane Fabrication

PLGA (poly(D,L-lactide-co-glycolide), lactide:glycolide = 50:50) pellets were purchased from Sigma-Aldrich (P2191). A 9 wt% PLGA solution was prepared by dissolving PLGA completely in acetone at 45 °C with magnetic stirring for 3 hours. Camphor (100 wt% relative to PLGA, 96%, Sigma-Aldrich) was then added to the PLGA solution and stirred at 45 °C for 1 hour. Before membrane fabrication, the PLGA–camphor solution was cooled to room temperature. PLGA membranes were fabricated by spin-coating the PLGA–camphor solution onto a silicon wafer at 400 rpm for 5 seconds. The coated wafer was subsequently immersed in distilled water at room temperature for 30 minutes to solidify the PLGA and camphor. To generate a porous membrane, camphor was removed by freeze-drying for 3 hours.

### Membrane Characterization

The morphology and internal porous architecture of the PLGA membranes were examined using scanning electron microscopy (SEM; Quanta 600 FEG, FEI Company, USA). To visualize cross-sectional features, the membranes were cryo-fractured in liquid nitrogen prior to imaging. Samples were fixed in 2.5% glutaraldehyde for 1 hour at room temperature, rinsed three times with phosphate-buffered saline (PBS), and subsequently post-fixed in 1% osmium tetroxide (OsO₄) for 1 hour at 4 °C. Following fixation, the samples were dehydrated through a graded ethanol series (50–100%) and further treated with hexamethyldisilazane (HDMS, Sigma-Aldrich) for 15 min before being air-dried.

Pore size distribution and surface porosity were quantified from SEM micrographs using ImageJ software. For each condition, pore diameters were measured from 100 randomly selected pores across four representative images, and porosity was calculated by threshold-based area analysis of four independent SEM fields.

### PLGA Membrane Degradation Assessment

To evaluate degradation, PLGA membranes were incubated in PBS at 37 °C for designated time periods. After incubation, the membranes were rinsed with distilled water and freeze-dried for 3 hours to remove residual moisture from the pores. The dry weight of the degraded membrane (W_t_) was then measured. The degradation ratio was calculated using the following equation:

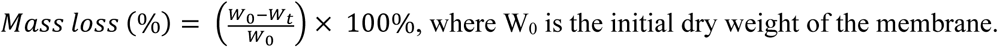

where W_0_ is the initial dry weight of the membrane.

### Permeability Measurement

To evaluate membrane permeability, the cell culture medium was first removed, and the device was gently rinsed with PBS. Subsequently, 1 mL of phenol red–free culture medium was introduced into the basolateral chamber, and 0.5 mL of FITC–dextran solution (4 kDa; Sigma-Aldrich) was added to the apical chamber. The chip was incubated at 37 °C, and aliquots were collected from the basolateral compartment at 0, 10, 20, 30, and 40 minutes into a black 96-well plate. Each sample was analyzed at the same time after collection to prevent signal loss. Fluorescence intensity was measured using a microplate reader (SpectraMax i3x) at excitation and emission wavelengths of 490 nm and 525 nm, respectively. The apparent permeability coefficient (P_app_, cm s^-1^) was determined using the following equation:

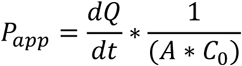

where dQ/dt represents the transport rate of FITC–dextran across the membrane, A is the effective membrane area (cm^2^), and C_0_ is the initial concentration of the fluorescent tracer in the apical compartment (mg mL^-1^).

### Mechanical Characterization

The mechanical behavior of the PLGA membranes was assessed through uniaxial tensile testing. Samples were stretched at a crosshead speed of 10 mm/min using a universal testing machine (MTS Criterion® 42) equipped with a 50 N load cell. Stress–strain responses were continuously recorded, and the apparent Young’s modulus was derived from the initial linear region of the stress–strain curves. Membrane thickness was determined from cross-sectional SEM images to ensure accurate normalization of stress values.

### Fourier Transform Infrared Spectroscopy

Fourier transform infrared (FT-IR) spectroscopy (PerkinElmer Inc., USA) was employed to evaluate the chemical structure of the PLGA membranes. Spectra of pure PLGA, camphor, and PLGA membranes were collected within the 600–2200 cm^-1^ range, using 32 scans per specimen to ensure sufficient signal resolution.

### Device Assembly

To fabricate the lung alveoli-on-a-chip device, a porous PLGA membrane was first bonded to the apical PDMS chamber (surface area: 1.13 cm²) using uncured PDMS as an adhesive, followed by curing at 40 °C overnight to ensure stable bonding. The membrane-attached apical chamber was then attached to the basolateral PDMS chamber (area: 2.01 cm^2^) using a thin layer of liquid PDMS, followed by overnight curing at 40 °C to ensure complete bonding.

### Cell culture

For cell culture, the lung-on-a-chip device was sterilized under UV light for 2 hours after being sprayed with 70% ethanol. Before seeding primary cells, the membrane was coated with 0.1% gelatin solution (Cell Biologics) at 37 °C for 5 minutes. To seed endothelial cells, the device was inverted, and either human lung microvascular endothelial cells (HMVECs, Lonza, P2–3) or human umbilical vein endothelial cells (HUVECs, ATCC) were seeded at a density of 2 × 10^5^ cells cm^-2^. After 3 hours of incubation, the device was returned to the upright position, and human primary alveolar epithelial cells (HPAECs, Cell Biologics, P2–3) were seeded in the apical chamber at a density of 5 × 10^5^ cells cm^-2^. Co-cultures of HPAECs and HMVECs were maintained in endothelial growth medium (EGM-2, Lonza, CC-3202) under liquid–liquid culture (LLC) conditions at 37 °C with 5% CO_2_ for up to 7 days. On day 7, the medium in the apical chamber was removed to initiate air–liquid interface (ALI) culture, which was maintained for 4 days before further experimentation. During ALI culture, the medium in the basolateral chamber was replaced every 2 days. Similarly, co-cultures of HPAECs and HUVECs were maintained in EGM-2 medium (Lonza, CC-3162) under identical conditions.

### Cell Viability Assessment

Cell viability was assessed using a Live/Dead Cell Viability and Cytotoxicity Assay Kit (Biotium, USA). Following gentle rinsing with PBS, samples were incubated at 37 °C for 30 min in a staining solution containing 2 µM calcein AM and 4 µM ethidium homodimer III (EthD-III) prepared in culture medium. After staining, the samples were washed twice with PBS and transferred to fresh medium prior to imaging. Fluorescence images were acquired using a Leica DMi8 microscope at 20× magnification, and cell viability was quantified by analyzing four randomly selected fields per sample.

### Immunofluorescence microscopy

The cells cultured on the membrane were washed three times with PBS and subsequently fixed in 4% paraformaldehyde (Sigma-Aldrich) for 15 minutes at room temperature. The membranes were then cut, and the fixed cells were permeabilized with 0.1% Triton X-100 in PBS for 10 minutes at room temperature. Following permeabilization, the samples were blocked with PBS containing 5% donkey serum for 45 minutes at room temperature. Primary antibodies, including ZO-1 (Rockland, 600-401-GU7, 1:400), VE-cadherin (Cell Signaling Technology, 2500, 1:200), laminin (Proteintech, 23498-1-AP, 1:200), collagen IV (Proteintech, 19674-1-AP, 1:200) and phospho-histone H2A.X (Cell Signaling Technology, 1:200), were diluted in blocking buffer and incubated with the samples overnight at 4 °C. After washing with PBS, the cells were incubated with secondary antibodies for 1 hour at room temperature. Nuclei were counterstained with DAPI following secondary antibody incubation. Fluorescence images were acquired using a Leica DMi8 fluorescence microscope.

### Transepithelial Electrical Resistance Measurement

Transepithelial electrical resistance (TEER) was measured to evaluate barrier integrity of the cell layers cultured on the PLGA membrane. Measurements were performed using a chopstick electrode system (MERSSTX01, Millipore, USA). The resistance of the cell layer (R_cell_) was obtained by subtracting the background resistance of the blank device (without cells) from the total measured value. TEER (Ω·cm^2^) was then calculated as the product of R_cell_ (Ω) and the effective membrane area (cm^2^). For ALI conditions, 300 µL of culture medium was added to the apical chamber immediately prior to measurement to ensure consistent electrical contact.

### Fluorescent PLGA Membrane Fabrication

To fabricate fluorescent PLGA membranes, FITC-labeled PLGA particles (Mw 5 kDa, Nanosoft Polymers) were added to the PLGA solution described above at a mass ratio of 1:20 (fluorescent PLGA : non-fluorescent PLGA).

### Fluorescence Image Quantification

For quantitative fluorescence analysis, four representative fields were captured at 20× magnification using a Leica DMi8 fluorescence microscope. The total fluorescence intensities of FITC-labeled PLGA, collagen IV, and laminin were measured in each image using ImageJ software (NIH, USA).

### Quantification of γ-H2AX–Positive Cells

The proportion of γ-H2AX–positive cells was determined by dividing the number of γ-H2AX–stained nuclei by the total number of nuclei within a defined area. For each sample, four independent fluorescence images were acquired and averaged to calculate the mean γ-H2AX positivity.

### Hydrogen Peroxide Quantification

Hydrogen peroxide (H_2_O_2_) levels were quantified using a Hydrogen Peroxide Assay Kit (Sigma-Aldrich, USA) following the manufacturer’s instructions.

### Statistics

Statistical analyses were conducted using Student’s t-test for pairwise comparisons. For experiments involving three or more groups, one-way or two-way analysis of variance (ANOVA) was applied, followed by Tukey’s post hoc test to assess pairwise differences. A p-value of less than 0.05 was considered statistically significant.

## Supporting information

upplemental File

