## Supplementary material for "A biodegradable porous membrane-based lung alveoli-on-a-chip for assessing particulate-matter-induced pulmonary toxicity": upplemental File

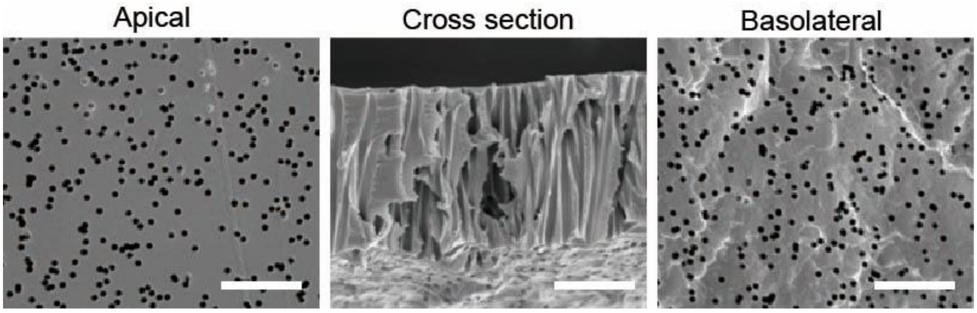


**Figure S1**. Representative SEM images of the Transwell® membrane from the apical surface, cross section, and basolateral surface. Scale bars: 5 μm.

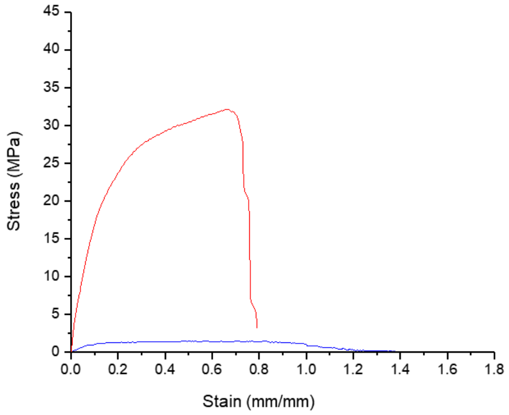


**Figure S2**. Tensile stress–strain curves of the Transwell® membrane (red) and the porous PLGA membrane before degradation (blue).

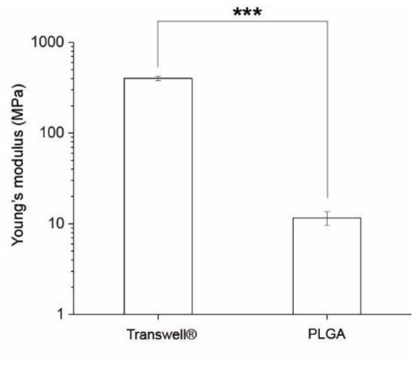


**Figure S3**. Young's modulus of the Transwell® membrane and the porous PLGA membrane before degradation (n = 4).

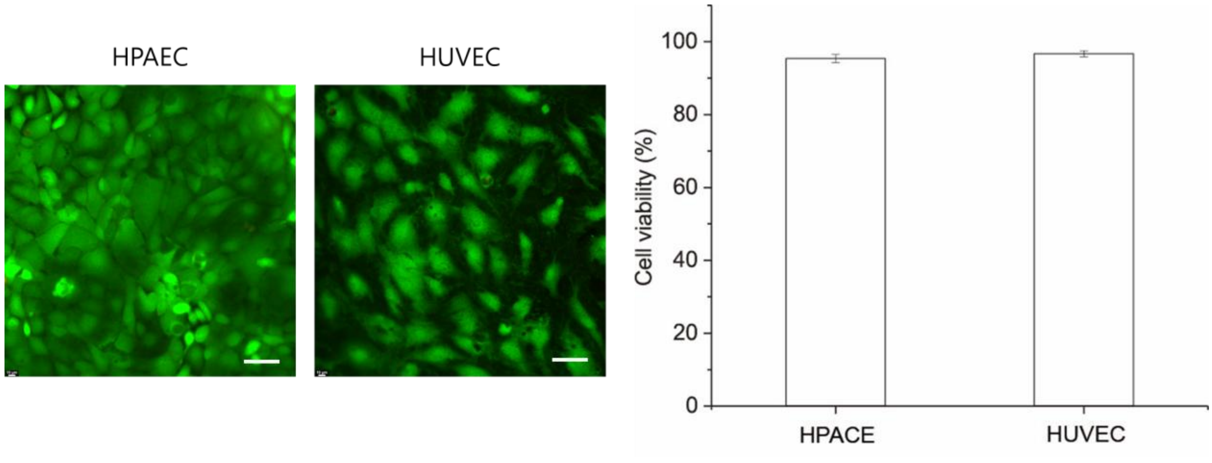


**Figure S4**. Live/dead staining images of co-cultured HPAECs and HUVECs in the chip after 11 days (left). Scale bars: 50 μm. Quantification of cell viability in the chip system after 11 days of HPAEC–HUVEC co-culture (right).

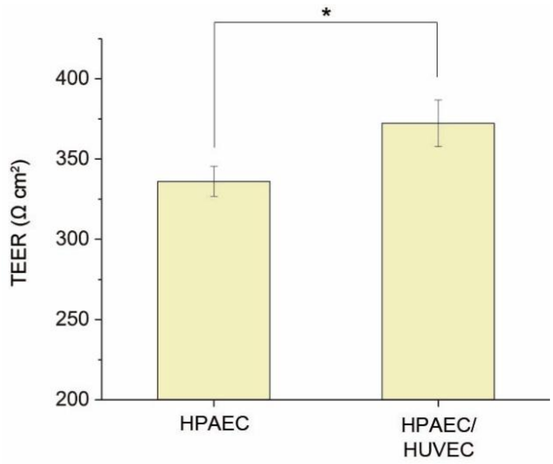


**Figure S5**. Quantification of transepithelial electrical resistance (TEER) in the chip system for HPAEC monoculture and HPAEC–HUVEC co-culture (n = 4).

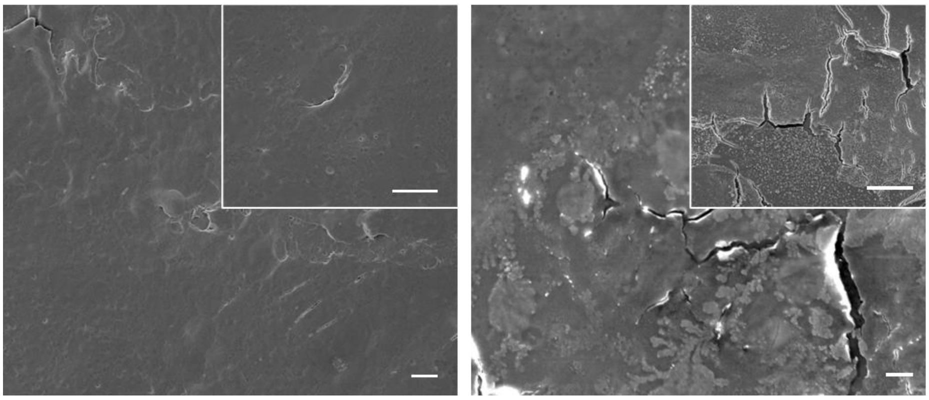

**Figure S6**. Representative top-view SEM images of cells cultured on PLGA membranes after 11 days of co-culture: HPAECs (left) and HUVECs (right). Insets show higher-magnification images. Scale bars: 10 μm.

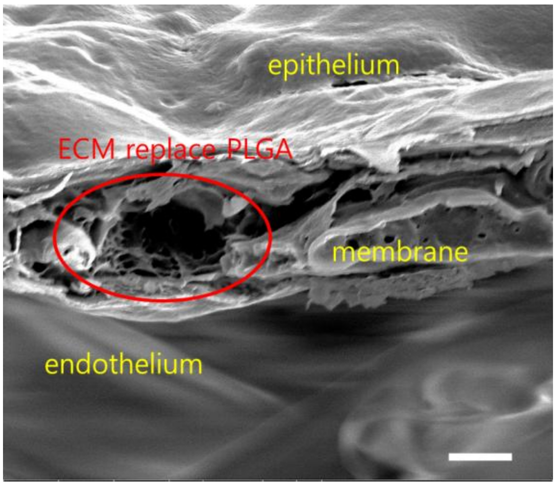


**Figure S7**, Representative cross-sectional SEM image of cells cultured on porous PLGA membranes under HPAEC–HUVEC co-culture conditions. The medium in the apical chamber was removed to expose the epithelial cells to air, establishing an ALI culture. Scale bar: 2 µm.

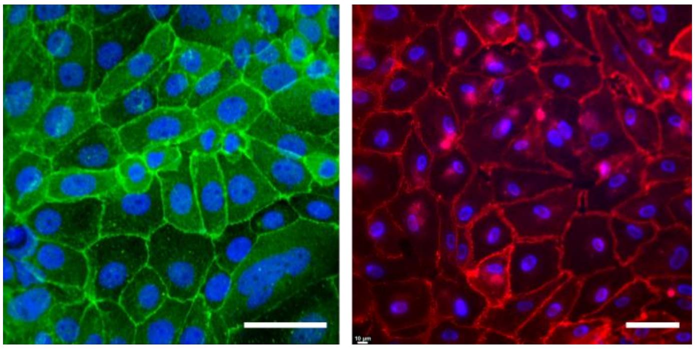

**Figure S8**. Representative immunofluorescence images of cells co-cultured on PLGA membranes: HPAECs stained for ZO-1 (left) and HUVECs stained for VE-cadherin (right). Nuclei are counterstained with DAPI (blue). Scale bar: 50 μm.

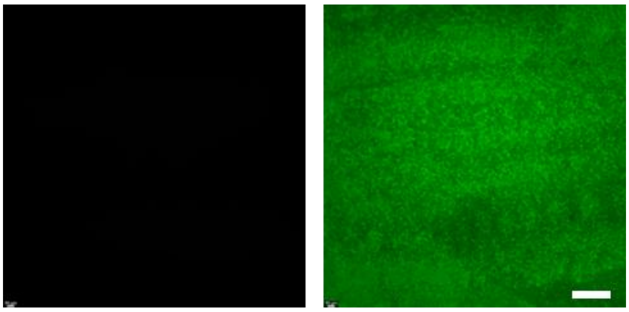

**Figure S9**. Fluorescence images of porous PLGA membranes without (left) and with (right) FITC–PLGA incorporation. Scale bar: 50 μm.

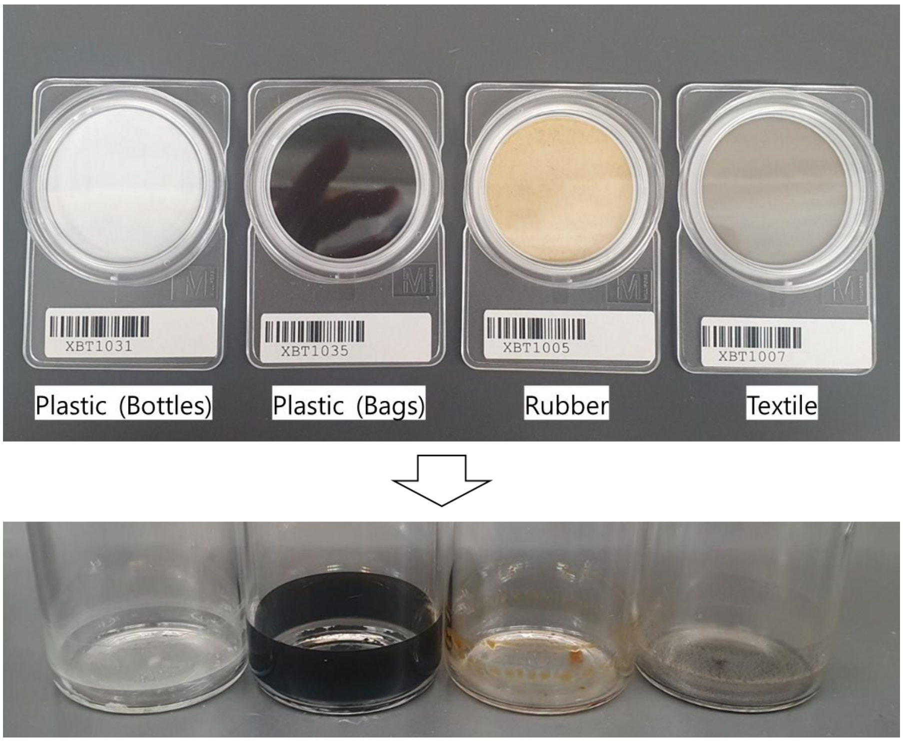


**Figure S10**. Four types of waste combustion products collected on membrane filters (top): plastic (bottles), plastic (bags), rubber, and textile. The corresponding particles separated from the filters are shown below (bottom).

| Species/ Waste Materials | Percent (%) of $PM_{2.5}$ Mass | | | |
|---|---|---|---|---|
| | Rubber | Textile | Plastic (Bottles) | Plastic (Bags) |
| HCl | 6.6908 | 1.0968 | 0.2162 | 0.2428 |
| S | 0.5772 | 0.2685 | 0.0000 | 0.0008 |
| Cl | 1.1251 | 0.5272 | 0.0420 | 0.0618 |
| Cu | 0.0289 | 0.0058 | 0.0009 | 0.0104 |
| Zn | 0.0198 | 0.0032 | 0.0000 | 0.0032 |
| Cd | 0.0005 | 0.0001 | 0.0001 | 0.0000 |
| Sb | 0.0489 | 0.0051 | 0.0000 | 0.0015 |
| Ce | 0.0116 | 0.0042 | 0.0052 | 0.0000 |
| Pb | 0.0198 | 0.0015 | 0.0019 | 0.0050 |
| Phthalates | 4.4445 | 0.1138 | 1.8912 | 1.6524 |
| PAHs | 1.0135 | 0.0315 | 1.3032 | 0.7100 |
| nitro-PAHs | 0.0067 | 0.0044 | 0.0082 | 0.0033 |

**Table S1**. Abudance (% of $PM_{2.5}$ mass) of major toxic chemicals from four types of waste combustion products.
